# Enhanced Prediction of Gut Microbiome–Related Diseases Using Hybrid Machine Learning Models

**DOI:** 10.64898/2026.06.24.734177

**Authors:** Sathya Abhijna Marisetti, Prathit Chatterjee, U. Deva Priyakumar

## Abstract

The human gut, containing 100 trillion microbes, is also considered the “second brain,” having control over the different functions of the physiological system. With advancements in bioinformatics and the development of sequencing technologies, researchers are able to explore the diversity and functional implications of gut microbiota (GM), which have become strongly associated with a variety of diseases. Microbial imbalance, or dysbiosis, acts as a biomarker for early detection and prognosis of a disease. Artificial Intelligence and Machine Learning (AI/ML) methods, although extensively used in predicting GM associated diseases, are seldom translated to having practical real-world outcomes, necessitating the design of robust AI/ML models applicable in real-world scenario. We have therefore come up with designing stacking-based ensemble architectures (EM1 and EM2), developed by integrating multiple ML-based learning algorithms for improving disease prediction accuracy. The GM datasets, after split into training and test sets, were eventually fed into the proposed two-layer ensemble models, which combines the output from standardized base learners via a meta-classifier, strengthening classification robustness as well as ensuring consistency in optimized performance across diverse datasets. Both the proposed hybrid ensemble models have emerged to be superior performers over all baseline and deep learning models, with an average accuracy of 0.87 and 0.84 respectively. By combining multiple learners, the proposed ensemble models outperform traditional single-algorithm-based approaches to attain higher accuracy and robustness on complex GM datasets.

**Key messages:**

- Development of stacking-based hybrid ensemble models (EM), which can be employed to integrate different AI/ML algorithms with better prediction accuracy of gut microbiome (GM)-associated diseases.
- Use of independent GM datasets with preprocessing methods such as SMOTE and PCA to address class imbalance and high dimensionality.
- All the proposed EM architectures are mostly superior to the existing state-of-the-art AI/ML methods (highest prediction accuracy: 0.87 and 0.84 with EM1 and EM2 models respectively) for GM diseases predictions.
- The cross-cohort validation demonstrates high prediction accuracy and robustness, (AUC values close to 0.98 and 0.99, for EM1 and EM2).
- These therefore demonstrate the effectiveness of EM frameworks for GM associated disease prediction, paving the way for corresponding applications in precision medicine.

## 1. Introduction

The human gut microbiome (GM) consists of billions of bacteria, viruses, fungi, and other microorganisms that work together to form a dynamic and complex arrangement in our digestive system. Because of its ability to regulate the many physiological system functions, the human gut, which is home to 100 trillion bacteria, is sometimes referred to as the “second brain” (Figure 1).[1] Essentially physiological brain consists of 500 million neurons, the total number decreased by a factor of 10^5^ in comparison to gut microbiota. These microorganisms are essential for maintaining the immune system, aiding digestion, and producing essential vitamins – all of which have a major impact on our overall health.[2] Recent advances in bioinformatics and sequencing technology have highlighted the enormous diversity and functions of GM, highlighting the major impact of the same on human health and diseases. It is also a useful tool for predicting diseases because of its association with a number of health problems. Dysbiosis,[3] which refers to changes in the composition and functions of GM, has been associated with various diseases, including autoimmune diseases, gastrointestinal diseases, metabolic disorders, and mental health issues. By analyzing the abundance profiles (real values of bacterial composition) and gene expression of microbial species, researchers can find biomarkers associated with certain diseases, which eventually serve as early warning indicators and can mitigate disease prognosis, improve therapeutic diagnosis and patient outcomes. Furthermore, marker profiles of GM indicates the presence and absence of bacteria.[4]

**Figure 1.**
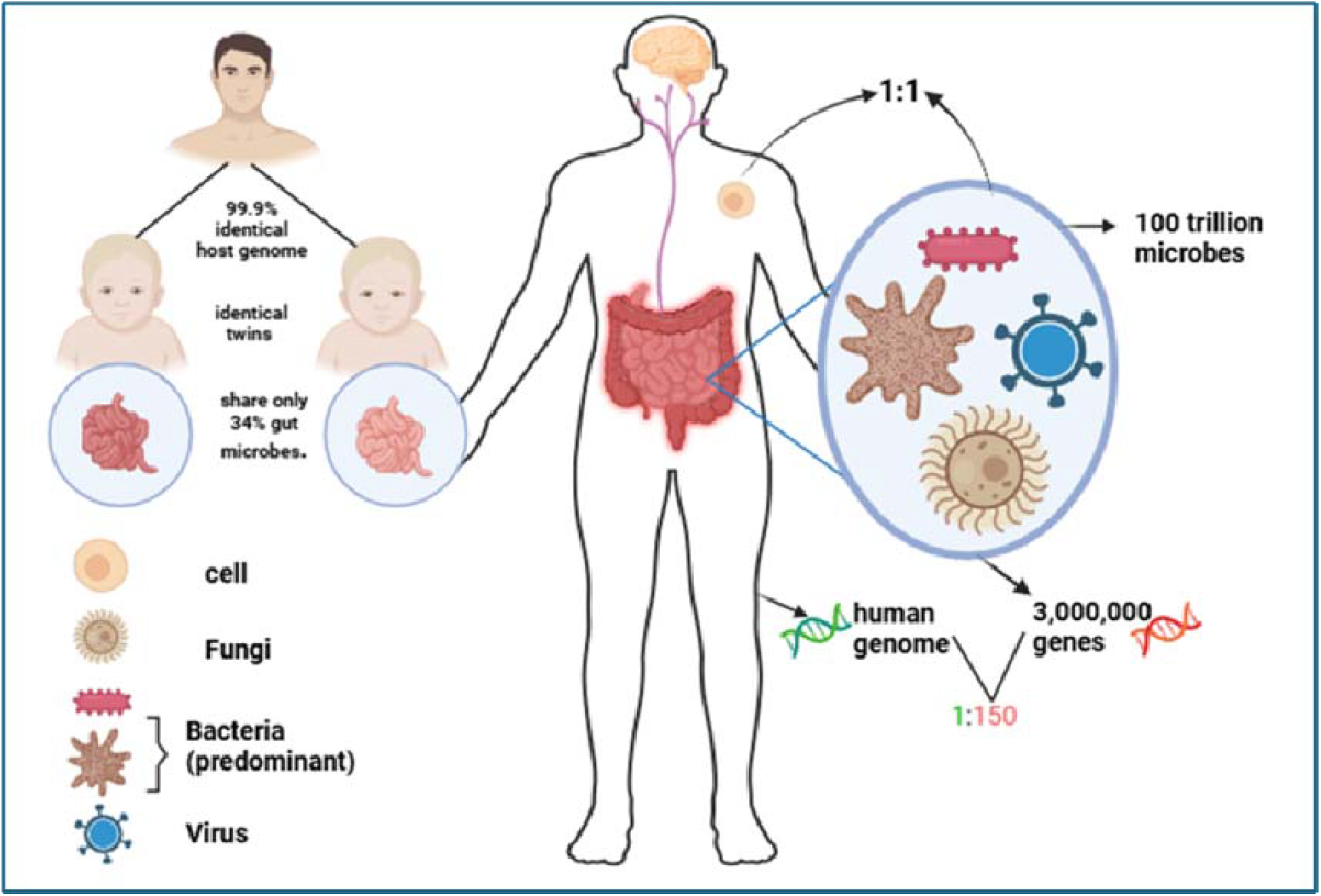
The human gut ecosystem

Both the abundance and marker profiles offer important information related to diseased conditions.[5] Quantitative profiling of microbial communities reveals disease-associated shifts in pathogenic and beneficial taxa, with functional profiles elucidating related metabolic and gene expression changes. [6, 7] Studies presenting the aforementioned factors associated with gut have been extensively undertaken, which reveal movement of microbes to and from the gut.[8] Additionally, the acellular transport of microbial antigens, influences physiological and pathophysiological processes that drive gut microbial ecology and can promote stomach diseases and cancer.[9] In 182 cirrhosis patients, particularly with acute-on-chronic liver failure, reduced gut microbial diversity and increased pathogenic taxa (Enterococcus, Pepto streptococcus) were associated with poorer survival and enriched ethanol, GABA, and endotoxin pathways.[10] Host genetics and gut microbiome interact in complex diseases, with specific taxa linked to genetic background and bidirectionally associated with multiple conditions including AF, CKD, and cancers.[11]. In order to reduce microbiome complexity and clarify the role of gut microbes in human health and disease, a guild-based approach has been used, in which functionally coherent bacteria are grouped.[12]

Previous studies underscored the importance of synthetic datasets, mathematical and statistical model analyses protocols, computational techniques, along with predictive Artificial Intelligence and Machine Learning (AI/ML) models.[13–16] These have been further useful in elucidating the relation between gut profiles and pathogenic diseases.[17–19] Predictive models utilise statistical methods on historical data to predict future events. therefore, they enable data-driven decisions, optimization of operations, and mitigation of risks. [20, 21] Integrating microbiome data into such models therefore allows not only the assessment of disease risk and progression but also enables precision medicine based on individual microbial profiles. It further facilitates biomarker discovery with accurate discrimination between healthy and diseased states for early detection, targeted therapies, and preventive healthcare.[22–24] Among the prevalent works, DeepMicro is a deep-learning framework using autoencoders, which transform high-dimensional gut-microbiome data into low-dimensional representations, enabling faster training and improved disease prediction on known datasets.[25] A Boosting GraphSAGE framework with metagenomic-disease graphs and DP-Net, helps predicting metabolic versus healthy samples, reaching high AUCs for IBD of 93% and colon cancer of 90%, outperforming prior ML/DL methods [26] MetaDR, a feature-fusion deep-learning model integrating known and unknown microbial abundances while preserving taxonomic relationships, enables interpretable, efficient, and biologically relevant disease predictions in metagenomics.[27] LogitBoost, a multi-class framework with backward feature elimination on microbial abundances across five taxonomic levels, effectively discriminated six disease categories, demonstrating its potential for microbial signature-based diagnosis. [28] Models like Support Vector Machines (SVM), Random Forest (RF), Artificial Neural Network (ANN), yielded 90% accuracy in distinguishing autism spectrum disorder (ASD) individuals from neurotypical ones, while pointing out key microbial predictors and how lesser-known gut microbes have been implicated in ASD.[29] TaxoNN, an ensemble neural network trained on stratified OTU clusters-scales better and outperforms classical methods with mean AUCs of 0.75 (cirrhosis), 0.92 (T2D), and 0.88.[30] In addition, ML models (RF, NN (Neural Network), SVM) identified key bacterial families, with RF achieving the highest performance (AUC 0.80) for predicting Parkinson’s Disease.[31] Several studies have reported reduced gut microbial alpha diversity in Parkinson’s disease and elevated diversity in anorexia nervosa, while predictive AUCs are 0.77, 0.74, and 0.74 for Alzheimer’s, multiple sclerosis, and schizophrenia, respectively.[32] RF, SVM, DT (Decision Tree), elastic net, NN were applied to the gut-microbiome features, yielding an AUC of 0.70 in cardiovascular disease prediction for the top 20 features.[33] GM Health Index (GMHI), based on arithmetic and mathematical principles, with microbial species as health indicators, yielded an overall accuracy of 73.7% in distinguishing between healthy and unhealthy subjects.[34] Statistics of 24-hr gut bacterial patterns in T2D have also been fed into RF models, successfully classifying the disease based on microbial signatures.[35]

ML ensemble models (EM) aggregate several weak learners to form a more powerful predictive algorithm, enhancing accuracy and stability.[36] A metaclassifier in ensemble models combines predictions from base classifiers to make a final decision, typically using models like logistic regression (LoR) or DT, chosen based on how well these learn from the base models’ outputs. Some of the most widely used ensemble methods are Bagging, Boosting, and Stacking, each intended to minimize variance, bias, or both, and maximize generalization. Ensemble learning is used in different fields like healthcare, finance, remote-sensing, traffic management, to name a few. [37–40] In recent times, EM’s have proven to be very effective in predicting disease from GM data, taking advantage of the strengths of several algorithms and compensating for individual weaknesses. EMs (including RF, Gradient Boosting, and stacked models) combine the predictions of several base models to improve robustness and accuracy. Diverse algorithms are combined in ensemble models, making them better at handling non-linear patterns, reducing overfitting, and improving generalization to unseen data.[41, 42]

Nevertheless, the domains on which EM architectures have been applied are comparatively much less than the overall fields of interest in healthcare. GM datasets (Table 1: Kaggle, Deepmicro, TaxoNN, Borenstein Lab), [25, 30, 43, 44] are usually complex, high-dimensional, and noisy, showing intricate interactions among microbial species that would influence the outcome of the disease. Traditional models may not capture these complex relationships well, leading to a deficiency in bridging the real-world outcomes with model performances. EMs can be effective at picking out weak microbial signatures linked to diseases that could be predicted with greater confidence and accuracy. In our current study, we have systematically constructed EM architectures, based on baseline model performances to elucidate diseased classifications from multiple gut-datasets. From our investigations, we present our findings and their implications, which are interpreted in the context of validating EM models, in retrieving key trends and patterns. Hybrid EMs have been found to surpass traditional ML algorithms and basic NN architectures (Table SI 6), leading to rational design and implementation of hybrid, multi-domain bioinformatics pipelines, while dealing with real-world GM data.

**Table 1.**
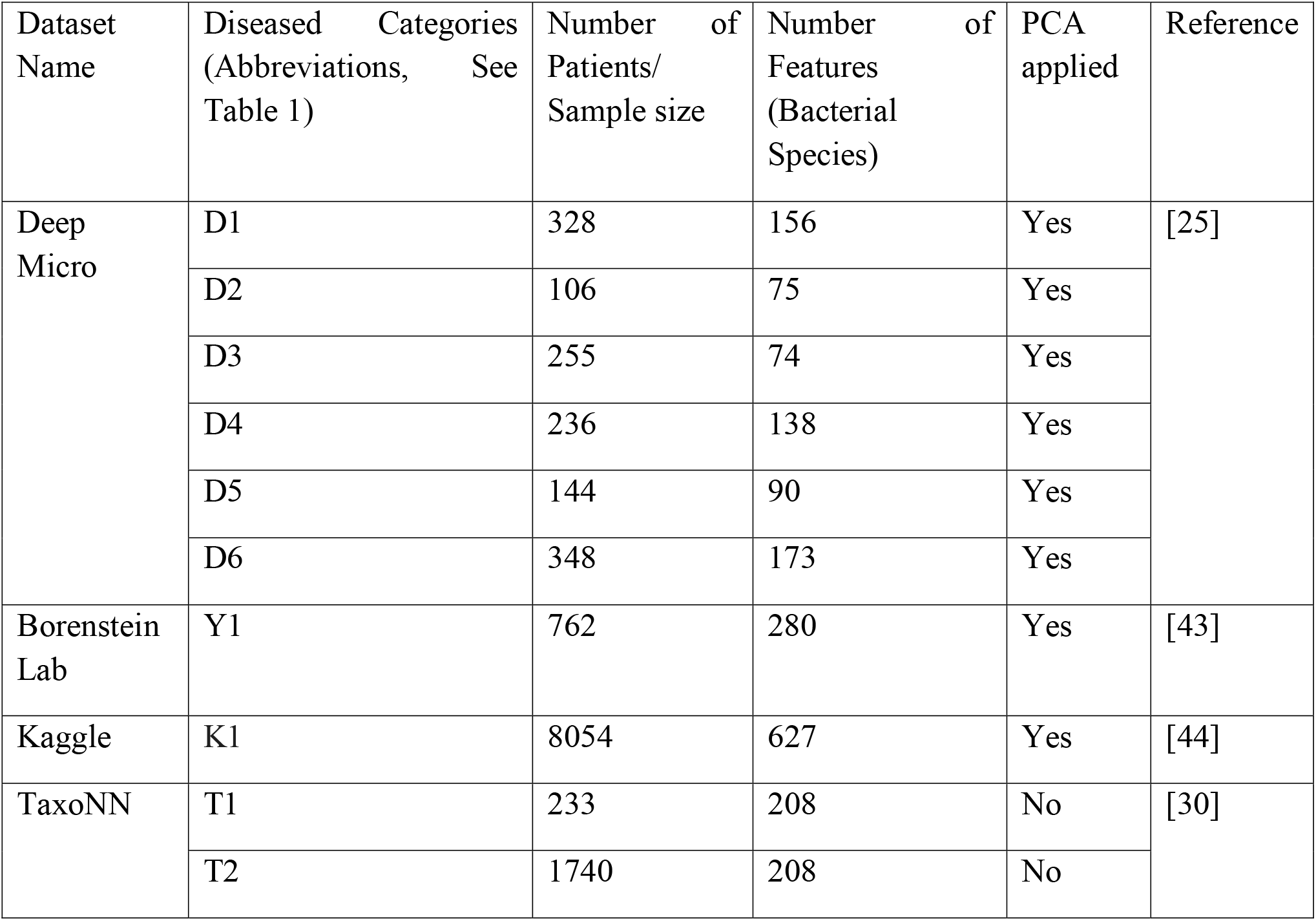
Dataset size after applying PCA. PCA was not applied to the Cirrhosis and T2D components of the TaxoNN dataset (T1 and T2, check abbreviations in SI (Table SI 1)), as the dimensions were already less (Table SI 3).

## 2. MATERIALS AND METHODS

### 2.1 Datasets

The data used in this work is drawn from multiple datasets (DeepMicro (D1 – D6), MetaML (K1), Borenstein Lab Dataset (Y1), TaxoNN Dataset (cirrhosis (T1), T2D (T2)).[25, 30, 43, 44] Each dataset contains the microbiome’s abundance profiles, and the target disease attributes along with metadata (corresponding original data dimension provided in Table SI 1). Each sample corresponding to each disease class (Table SI 1) is represented by a vector of GM abundance values. Major features in this work are taxonomic abundance profiles obtained from metagenomic sequencing, whereby each feature represents the relative abundance of a microbial taxon in a particular sample (the corresponding dimensions in Table SI 2). These profiles depict the functional and compositional changes of the gut microbiome linked to the health status of the host, disease course, inflammation, and metabolic dysregulation. In addition, the abundance features capture the presence–absence patterns as well as the quantitative changes in the populations of the microbiota, thus allowing the model to learn both disease-associated GM signatures and community-changes dependent on disease progression.

### 2.2 Data pre-processing

#### 2.2.1 Data cleaning

Data cleaning includes removing unwanted attributes (not contributing to the model’s learning process, removing the Nan (not-another-number)-containing values, and the attributes containing entire column as Nan). This is followed by filling the null values with the average of the given attributes in some cases.

#### 2.2.2 Data normalization

We apply a min-max scaler to bring uniformity and all the numerical values of a given attribute between 0 and 1. Using the min–max scaler increases the model’s performance and ensures consistent model behavior (eqn. 2):

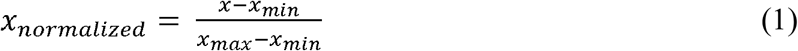

here, *x*_*normalized*_ is the normalized value scaled between 0 and 1, *x* the original value, *x*_*min*_ is the minimum value, and *x*_*max*_, the corresponding maximum value in the dataset.

#### 2.2.3 Data augmentation

To reduce the imbalance in the classes and avoid overfitting or underfitting, we have used SMOTE (Synthetic Minority Over–Sampling Technique) on D1, D2, D3, D4, D5 and D6, the smaller dimension datasets, as well as on the larger dimension datasets Y1, K1, and T2 (Table SI 2), in achieving a more balanced distribution across all the classes (the dimensions after augmentation provided in Table SI 3). The SMOTE algorithm was implemented for data augmentation, and created new instances, rather than duplicating minority classes.[45] It generates synthetic data for the minority class with fewer samples in the dataset. This algorithm initially identifies the minority class. For each minority sample *x*_*i*_, the k-nearest neighbor from the same class is selected, followed by selection of a random nearest neighbor *x*_*j*_, in eventually obtaining the midpoint of *x*_*i*_ and *x*_*j*_ to be the new synthetic data point (eqn. 1). The SMOTE algorithm is used mainly for augmenting the data for hybrid models. For the Kaggle dataset, we experimented with implementing the augmentation:

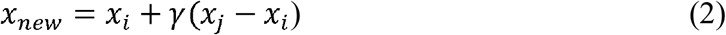

Here *x*_*new*_ is the synthetic data point that is generated. *x*_*i*_ is the data point from the minority class, γ is the random number between 0 and 1, and *x*_*j*_ is the randomly selected nearest neighbor of *x*_*i*_ (from the k-nearest neighbors).

#### 2.2.4 Data dimensionality reduction

This technique reduces dataset features while keeping essential information. We perform dimensionality reduction in order to improve the performance of the model and reduce the computational complexity. We have used principal component analysis (PCA: section 2.2.4.1) in undertaking the dimensionality reduction.

##### 2.2.4.1 Principal component analysis

PCA is the most popularly applied dimensionality reduction technique in the domains of data analysis and machine learning.[46] This transforms a dataset which could absolutely consist of time-dependent correlated features into a set of orthogonally transformed linearly uncorrelated variables or principal components, maximizing retention of variance within the data. Here, variance quantifies the proportion of data variability captured along each principal direction. Consequently, information loss is avoided, and simplification occurs. PCA is particularly helpful in dealing with high-dimensional data, to be visualizable in possible lower dimensions, thereby reducing overfitting and computational complexity, in overall improvement of the performance of ML algorithms. It was applied in order to retain the minimum number of principal components explaining 95% of the cumulative variance, hence preserving most of the informative structure of the data while reducing dimensionality and noise.

The key steps in PCA are standardizing the data, calculating the covariance matrix to understand the relationships between features, and finally obtaining the eigenvalues and eigenvectors of the covariance matrix. The eigenvectors provide a representation of the orientations of the transformed feature space, whereas their corresponding eigenvalues denote their magnitudes. After ordering the eigenvalues and choosing the leading (*k*) principal components, the original dataset is projected onto newly obtained coordinate systems, representing it in reduced dimensions. Here, it retains the most relevant characteristics of data but discards the least contributing aspects of its variability. After PCA, the final dataset obtained (Table 1) is split into a test set and a train set in the ratio of 80:20 and evaluated with different models (section 2.4).

Given a dataset X with n samples and m features, each feature is mean-centred (eqn. 3) to 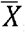 ,

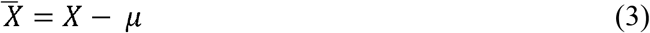

Here, *μ* (eqn. 3) is the mean vector of the dataset:

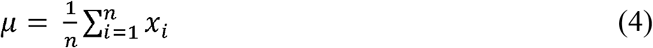

The covariance matrix C of the centred dataset is given in (eqn. 5),

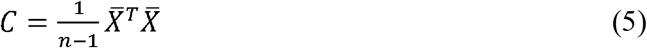

Finally, we solve the eigenvalue,

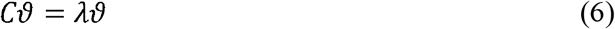

Here, *ϑ* are the eigenvectors and *λ* is the eigenvalue in (eqn. 6). Generally, the top *k* eigenvectors are selected corresponding to the largest *k* eigenvalues to form the transformation matrix *W*_*k*_ as shown here (eqn. 7):

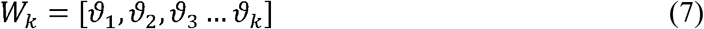

The dataset is transformed into the new lower-dimensional space as,

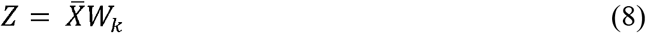

*Z* (eqn. 8) is the reduced representation (Table 1) of the originally augmented dataset (Table SI 3).

## 2.3 ML based models

ML based models are types of algorithms that learn patterns from data to make predictions or decisions without any explicit programming. They can be widely classified as supervised learning (e.g., linear regression (LR), LoR, SVM, DT, RF), unsupervised learning (e.g., k-means, PCA, DBSCAN), and reinforcement learning (RL) techniques (Q-learning, deep Q-networks).[47–49] Here, supervised learning models are been used owing to the datasets having the targeted variables, accumulated with the ground truths. A detailed discussion on the baseline models has been provided in Supplementary Information (Section SI 2).

### 2.4 Ensemble Models (EM)

Before implementing the EM Architecture (Figure 2), base models have been implemented on chosen GM datasets (Table SI 1, Table 1) like Random Forest (RF), Support Vector Machine (SVM), Extreme Gradient Boost (XGB), Naïve Bayes (NB), and Multi-layer perceptron (MLP) (Section SI 2). More accurate predictions are selected and tested with different combinations in the constructed EMs. The proposed approach utilizes stacking-based ensemble architecture to optimize the classification performance of this model, by combining different learning algorithms.[36] The input data is divided into training and test sets (80:20). In essence, it means that while training was done on 80% of each of these datasets, 20% is assigned as test. Using the training set, different combinations of the base models are fitted. With multiple trials with the EM variants, we narrowed down to two that performed quite strongly and uniformly well on all the six datasets (augmented and with reduced dimensions; Table 1). Keeping consistency with our experiments, the train-test split maintains a fixed random seed value. The training data undergoes preprocessing and reshaping, after splitting into training and test datasets. In general, the EM architecture makes use of a two-layer stacking ensemble technique to strengthen classification by gathering multiple base models with a meta-model. In the first layer, several base models are trained separately on flattened training data. A standard scaler is used for standardizing this flattened data. The outputs from different base models are combined into a single dataset and then been passed to the meta-classifier for final classification task. Additional information corresponding to each of these models are further explained below.

**Figure 2.**
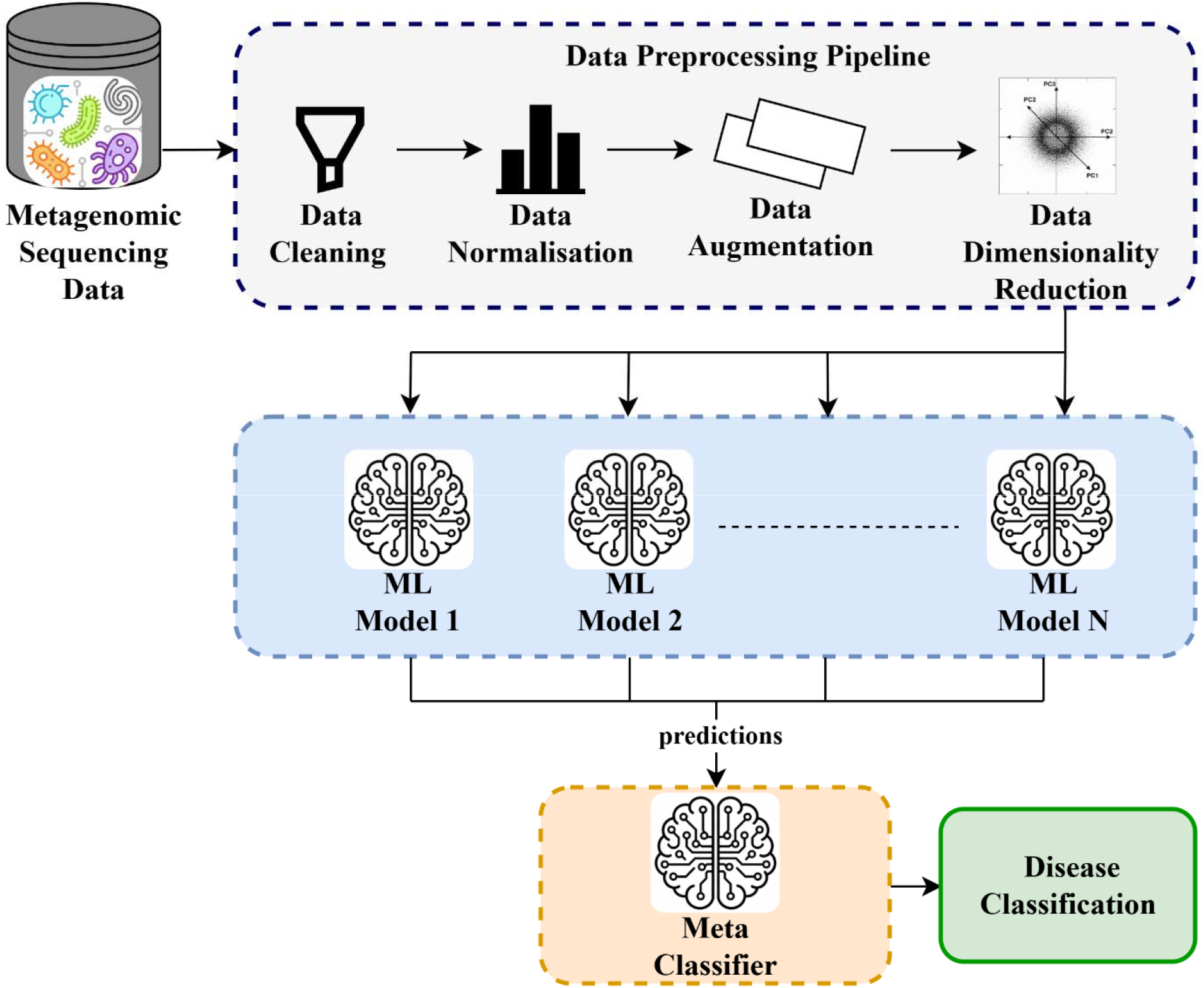
Ensemble models architecture

### 2.4.1 Ensemble Model-1 (EM1: NB+ XGB + SVM + RF + LoR)

For the current model, in the first layer, several base models for training separately on flattened training data include RF, SVM, XGB, NB, and LoR classifiers, each contributing its strengths to the ensemble. In this ensemble, the tree-based model RF is combined with a probabilistic linear classifier model (NB), a margin-based classifier (SVM), and XGB (Table SI 5). Meta-features are learned from the concatenated outputs of all base models and are used to train a LoR model, which will be the meta-learner that eventually integrates the predictions coming from the base models and generates the final output. Here, XGB and RF take the most complex relationship present in data, which could not be linearly correlated. The SVM works with more boundary separation optimality for good classification. NB and LoR contribute to diversity by handling probabilities more accurately, besides having linear separability. This combination of algorithms provides a more comprehensive approach to capturing various types of patterns in the data with high accuracy and generalization.

### 2.4.2 Ensemble Model-2 (EM2: RF +SVM + MLP)

In EM2, for each model, the features (as discussed before, section 2.4) are flattened before being fed into the models. This is because most of the models, especially SVM and MLP, might seem to be sensitive to feature scaling. In this ensemble model, the tree based model Random Forest, margin-based classifier SVM acts as the base models. (Table SI 5). MLP (a neural network, being the meta-learner) aggregates predictions from base models. This combination leverages the advantages of each model: RF’s robustness and resistance to overfitting, SVM’s precision and accuracy in boundary formation, and MLP’s capability to capture complex nonlinear relationships. When combined, they deliver balanced and efficient performance, especially in contexts where both linear and non-linear patterns are relevant, with each model addressing the shortcomings of the others.

### 2.4.3 Combinations with different Metaclassifiers

After observing that the EM1 and EM2 models outperformed the baseline models, we conducted further experiments using different combinations of meta-classifiers and baseline models inspired from EM1 and EM2 (Table SI 5). These have been implemented similar to previous protocol, using four (and two) models as base learners and one model as the meta-classifier, for EM1 (and EM2). A comparison of these models are discussed in section 3.1.

## 3. Results and Discussions

We present the findings of our investigation and discuss their implications in relation to validating GM-related ML models, highlighting key trends and patterns. Comparisons with previous literature are also drawn up to place the results into context. To compare the increase in performances of EM models over the baseline models (SI Section 2), we have used different metrics like Accuracy, Precision, Recall, F1-score, AUC, to evaluate the results for binary and multi-classification (SI Section 3).

### 3.1 Comparison between baseline and EM models

To compare the performances of the baseline and the ensemble algorithms, we hereby present the calculated Accuracy of the individual datasets when experimenting with different baseline and ensemble algorithms (Table 2). Among the different combinations, ensemble algorithms EM1 and EM2 (Section 2.4) were noticeably better than the base models. The average accuracy for ensemble models (EM1: 0.87, EM2: 0.84) are higher compared to any of the baseline models. The maximum increase in average accuracy from an ML model (KNN: 0.66 for K1, Table 2) to the ensemble algorithms is ∼ 32%. When we observed that the EM1 and EM2 models outperformed the baseline models, we further experimented with different variations of hybrid ensemble models (see Section 2.4.3). The majority of EM1 and EM2 variants, specifically EM1.1, EM1.2, EM1.3, EM1.4, as well as EM2.1, EM2.2, demonstrated superior performances (in terms of accuracy), compared to the baseline models (Table SI 6, Table 2). We however chose to introspect further on EM1 and EM2 as our specific case-study, by elucidating the performances for the imbalanced and balanced classes, the ML metrics, and comparing with the existing data of available literature.

**Table 2.** Average accuracy of individual datasets for hybrid models.

| Dataset | KNN | NB | RF | SVM | EM1 | EM2 | MLP | NN |
| --- | --- | --- | --- | --- | --- | --- | --- | --- |
| D1 | 0.60 | 0.66 | 0.69 | 0.86 | 0.89 | 0.80 | 0.87 | 0.63 |
| D2 | 0.72 | 0.63 | 0.50 | 0.72 | 0.68 | 0.63 | 0.81 | 0.63 |
| D3 | 0.96 | 0.88 | 0.96 | 0.96 | 0.98 | 0.80 | 0.78 | 0.41 |
| D4 | 0.83 | 0.70 | 0.87 | 0.93 | 0.94 | 0.92 | 0.91 | 0.91 |
| D5 | 0.37 | 0.55 | 0.65 | 0.82 | 0.65 | 0.69 | 0.55 | 0.44 |
| D6 | 0.70 | 0.71 | 0.62 | 0.92 | 0.88 | 0.86 | 0.81 | 0.71 |
| K1 | 0.66 | 0.31 | 0.63 | 0.67 | 0.95 | 0.98 | 0.68 | 0.66 |
| Y1 | 0.60 | 0.51 | 0.86 | 0.85 | 0.92 | 0.92 | 0.82 | 0.73 |
| T1 | 0.65 | 0.80 | 0.85 | 0.80 | 0.89 | 0.91 | 0.87 | 0.74 |
| T2 | 0.52 | 0.65 | 0.63 | 0.60 | 0.98 | 0.98 | 0.55 | 0.63 |
| Average accuracy | 0.66 | 0.64 | 0.72 | 0.81 | 0.87 | 0.84 | 0.76 | 0.64 |

### 3.2 Imbalanced and balanced class performances

Previously we observed the outperformances of the ensemble methods over baseline models (Table 2), where all the datasets have been already balanced (Table 1). To further verify the superiority of the EM models, and the relevance of data augmentation (see section 2.2.3) in addressing class imbalance, we calculated the conventional ML model metrics corresponding to the selected EM architectures (see section 3.1), for the unbalanced and balanced classes (Tables SI 7, SI 8). The datasets (K1, Y1, T1, T2) chosen for the aforementioned calculations were based on increased performances in Accuracies (Table 2), in spite of these being multi-class datasets with higher dimensionality (Table 1). Data augmentation was not applied to T1, as the class distribution was already balanced (see Table SI 2). Table SI 7 shows the metrics on the K1 (with 13 diseased classes), Y1, and T2 (with binary classes of whether that particular disease is present or not), before class balancing. In comparison, Table SI 8 shows the increased performance for all the chosen datasets once the classes are balanced.

From Figure 3, the improvement in performance metrics is shown for both EM1 and EM2 hybrid models after class balancing. It is calculated as the percentage change or the difference between the metric values after and before class imbalance, normalized by the metric value before class imbalance, and multiplied by 100. EM1 takes the lead here in improving Precision (+453%), F1-Score (+313%), and Recall (+191%), indicating that it is better tailored for the imbalanced data. There is some improvement for EM2 as well across all metrics, though not to the same extent as EM1, except for CV-Score (+86%), where EM2 performs slightly better than EM1. The results for both models for AUC are rather close (∼30%). Although EM1 outperforms classification performance, the improvements in ML metrics for the chosen EM models underscores that GM disease predictions are sensitive to class imbalance.

**Figure 3.**
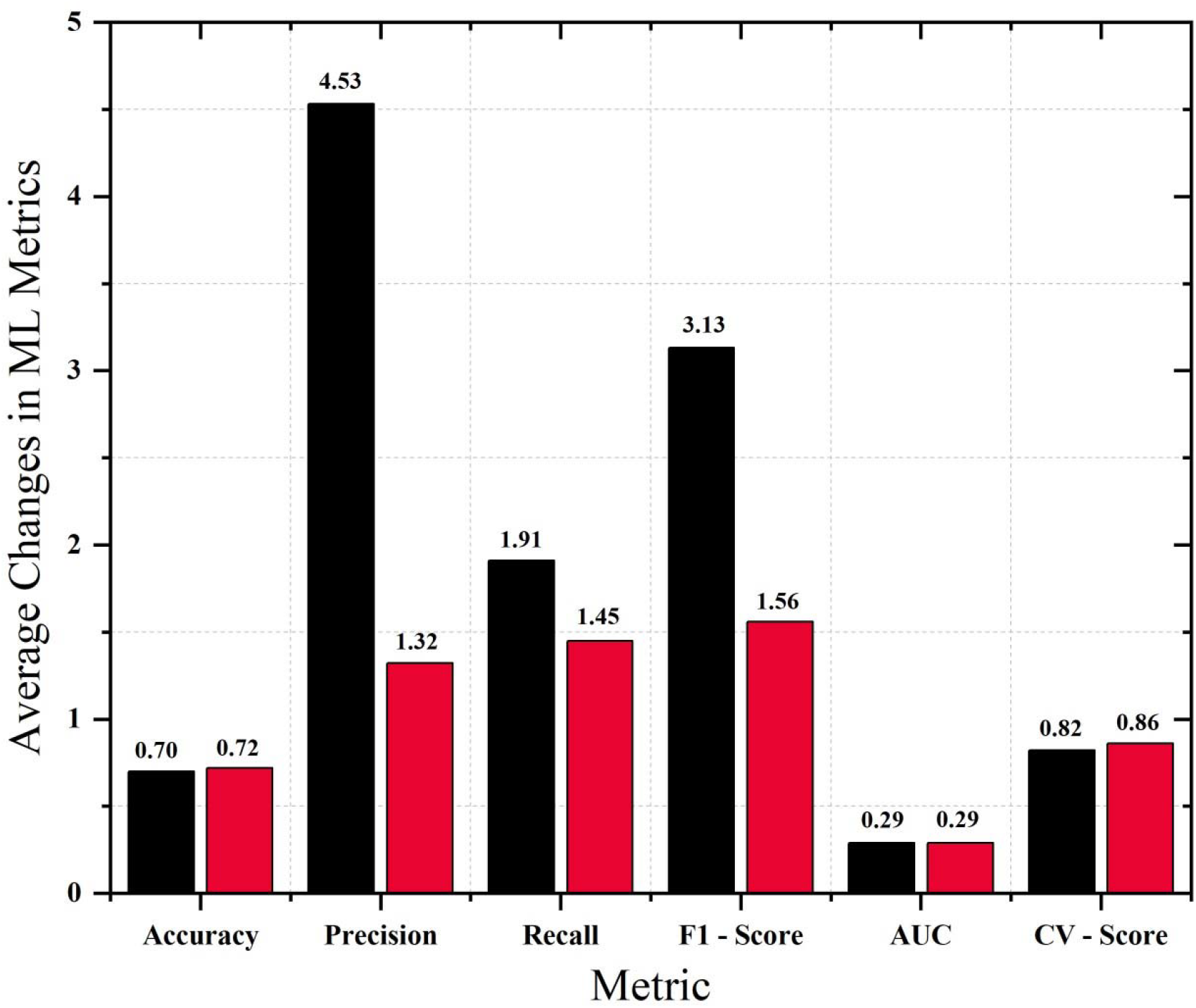
Increase in performance metrics from the imbalanced (Table SI 7) to balanced (Table SI 8) class performances, averaged over the selected datasets (K1, Y1, T2), obtained separately corresponding to the hybrid models (EM1 (*black*), EM2 (*red*)).

### 3.3 ML metrics comparisons

Initially, we evaluated multiple baseline ML models—including KNN, NB, RF, SVM, MLP, and standard NN - on the ten datasets (Table 2). As presented in Table 2, the average accuracy of EM1 and EM2 were 0.87 and 0.84, respectively, clearly outperforming all traditional models. For comparison, the average accuracy of the baseline models range between 0.61-0.81. Particularly, EM1 achieved top accuracy on datasets like D1 (0.89), D3 (0.98), D4 (0.94), and T2 (0.98), while EM2 excelled on K1 (0.98) and T1 (0.91), some of these were multi-class high-dimensional datasets (Table 1). These indicate not only more accurate predictive performances but also robustness across diverse biomedical domains for the constructed ensemble architectures.

In terms of elucidating the performance differences between imbalanced and balanced classes (Section 3.2), EM1 and EM2 have shown promising results in generalizability and resilience to class imbalance (Tables SI 7, SI 8). Before balancing, Y1 was highly imbalanced, and ensemble models showed lesser to moderate performances (Table SI 7: EM1-accuracy 0.43, AUC 0.71; EM2 - accuracy 0.41, AUC 0.69). However, after class balancing (Table SI 8), EM1’s accuracy surged to 0.92 (AUC 0.98) and EM2 to 0.92 (AUC 0.99). Similarly, in K1, accuracy improved from 70% to at least 95%, with AUC reaching 0.99 for both models. T2 showed near-perfect performances with EM1 (0.98 accuracy) and EM2 at (0.98 accuracy), both maintaining an AUC of 0.99. Even for T1, a balanced dataset, both models maintained high accuracy (0.89 and 0.91) respectively and AUC values are 0.95 and 0.92 respectively (Table SI 8).

Figure 4 gives a comparative assessment of the two ensemble models EM1 and EM2 on four datasets based on microbiome data, K1, Y1, T1, and T2. The models consistently demonstrated good performance across all datasets, although the maximum accuracy and F1 scores were obtained for T2 (0.98). EM1 performed slightly better than EM2 on K1, Y1, and T2, while T1 showed fairly comparable results between these two models. Cross-validation scores suggest good generalization (Figure 4 c, Table SI 8). To compare the performances of the datasets to those of the original studies, we have chosen the AUC metrics (Figure 4 d), as AUC have been mostly used in the previous works to validate the corresponding performances of the proposed models. From our study, AUC confirms the advantage for the proposed ensemble models over the models from the original study, especially on K1, Y1 and T2, where EM1 and EM2 achieve near perfect AUC scores (0.98-0.99) and outperformed the baseline from the original paper. In conclusion, the results confirm the strength and reliability of the proposed hybrid approaches across the chosen (diverse) datasets.

**Figure 4.**
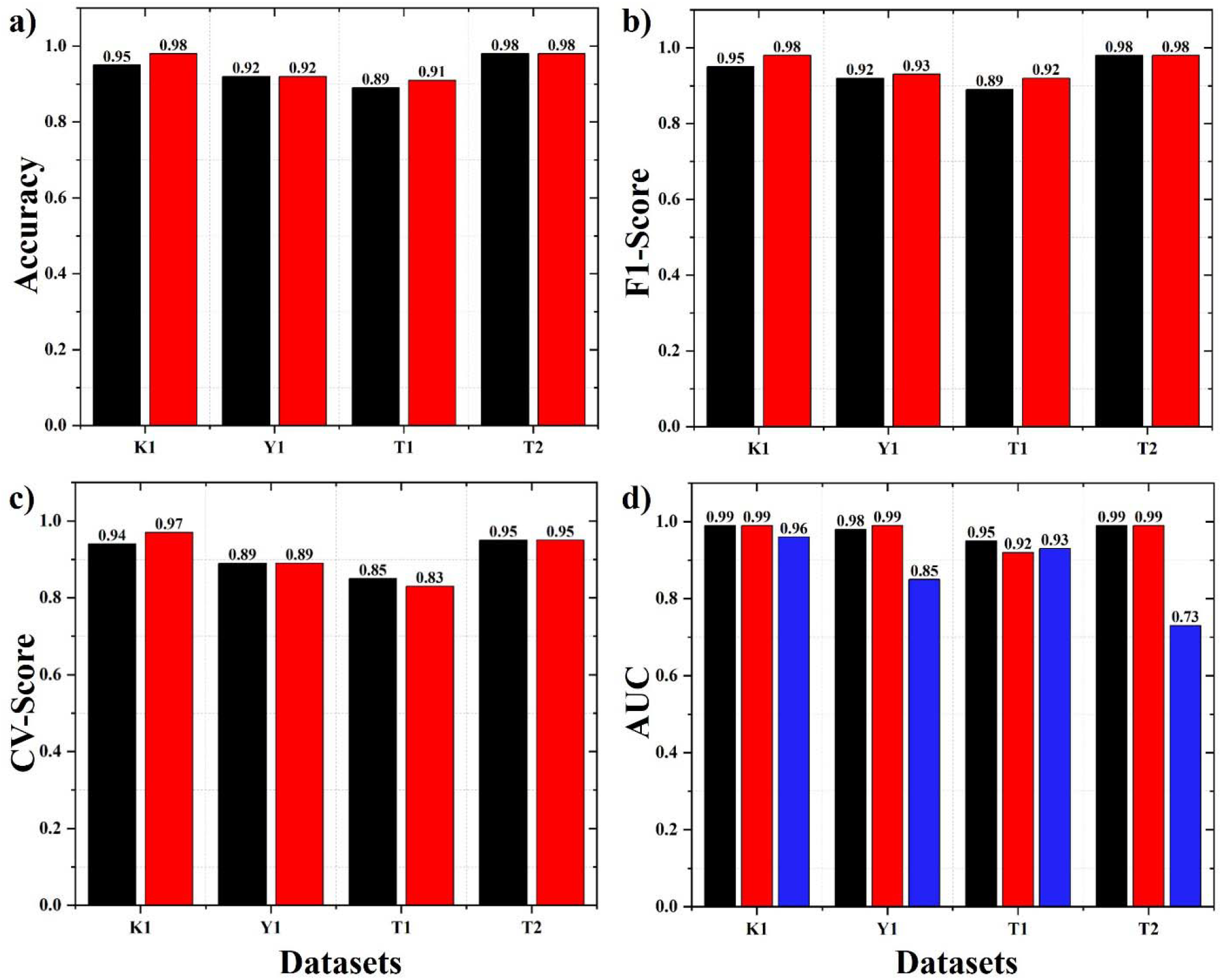
ML metrics for hybrid EM models (EM1 (*black*), EM2 (*red*)): a) Accuracy, b) F1 score, c) CV score; d) Comparison of AUC computed for the hybrid EM models with those of the original studies (*blue*).

Notably, both ensemble models consistently outperformed deep learning models (MLP and NN) used for our model benchmarking. While MLP and NN achieved average accuracy of 0.76 and 0.64, respectively, EM1 and EM2 had average accuracy of 0.87 and 0.84, respectively (Table 2). EM1 further outperformed NN on all individual datasets. Therefore, ensemble models exhibited lower variance in performance across datasets compared to neural networks, highlighting their greater stability and reliability, for GM associated features and dimensions.

In order to validate the constructed models for multi-class classification, we had a closer look at class-wise F1 scores across all datasets. Table SI 9 shows that both EM1 and EM2 generally have strong, reliable performance but vary significantly across disease categories. Indeed, for most of the datasets—DeepMicro (D3, D4), Borenstein Lab (Y1), Kaggle (K1), and TaxoNN (T1, T2)—the models produce consistently high F1 scores ≥0.90, reflecting good generalization to well-represented and biologically distinct phenotypes such as IBD Crohn’s disease, IBD ulcerative colitis, Multiple Polypoid, Large Adenoma, and Stec2-positive conditions. These classes exhibit clear microbial signatures and benefit from the models’ capacity to distinguish strong disease-specific patterns. However, in the case of subtle microbiome shifts or overlapping phenotypes of Normal, Normal at risk, and T2D, its performance is more variable, with F1 scores occasionally falling into the 0.55–0.75 range, e.g., D2-Normal, D5-Normal. Such borderline or physiologically diverse categories introduce more intra-class variability and are hence harder to classify reliably. When comparing EM1 and EM2, minor but consistent gains for EM2 can be seen in several datasets, including Y1 and K1, while EM1 performs slightly better for difficult classes in DeepMicro, such as D1 Obesity/Leanness. Overall, the models perform very well for well-defined disease phenotypes, whereas for more heterogeneous “Normal” or metabolic disorder classes, it has less discriminative power.

## 4. Conclusion

We conclude that a well-constructed hybrid model always proves to be much superior in the task of disease classification when compared with the existing established benchmark models and individual base learners on several datasets in terms of accuracy and generalizability, while overcoming the class imbalance. Combining several classifiers within a hybrid framework has greatly enhanced predictive accuracy as well as the ability of the model to discriminate between classes. Their continuous outperformance in both classical models and deep learning models indicates that they are well-suited for practical applications of real clinical data, which is problematic in terms of unevenness in distribution and high domain variability. It thus offers improved robustness and reliability, which are important factors for clinical decision making and medical diagnosis. Class-wise F1 analysis demonstrates that hybrid models excel at capturing strong disease-specific microbiome signatures, while highlighting the intrinsic difficulty of discriminating heterogeneous or subtly varying phenotypes. A major future enhancement could, however, be the integration of multi-omics data as well as explainable AI/ML in depicting predominant class features, where we could have an integrated and system-wide view of disease mechanisms. Personalized medicine approaches need to be designed on the microbiome profiles of individuals to advance effective interventions. Creating attention-driven architectures, such as Transformer models, for biomedical sequences will also be one of the best ways to capture considerably sophisticated patterns and contextual dependencies in the coming years. This will make disease prediction more accurate and reliable, supporting the rational development and validation of hybrid, multi-domain bioinformatics pipelines under real-world gut microbiome data constraints.

## Supporting information

Supplementary Word File containing the supporting information, including protocols, algorithms, methods, figures, and tables.

## Acknowledgment

SAM thanks Ms. Ananaya Jain for initial assistance with the data understanding. The authors thank IHub-Data, IIIT-H for funding and support. U.D.P. thanks DST-SERB (CRG/2021/008036) and Kohli Center on Intelligent Systems, IIIT Hyderabad for support.

## Author contribution

UDP and PC conceived and supervised the project. SAM undertook literature review, model constructions and analyses. SAM and PC wrote the manuscript. All authors read and approved the final version.

## Code Availability

The source code used for data analysis and model implementation in this study is available at https://github.com/devalab/ML_GUT.

## Data Availability

The data used for this GM-related case-study are available publicly. The datasets which have been used are Deepmicro, data available from Borenstein lab, from Kaggle, and TaxoNN. Details of the datasets (appropriately referenced) are provided in (Table 1, and SI Table 1).

## Acknowledgment

SAM thanks Ms. Ananaya Jain for initial assistance with the data understanding. The authors thank IHub-Data, IIIT-H for funding and support. U.D.P. thanks DST-SERB (CRG/2021/008036), IHub-Data, IIIT Hyderabad, and Kohli Center on Intelligent Systems, IIIT Hyderabad for support.

**U. Deva Priyakumar** is a Professor at IIIT Hyderabad, and his research group works on applying modern machine learning methods to problems in computational chemistry and biology. They also develop large-scale datasets that are useful for machine learning applications in drug discovery.

## References

[1] Shanahan F, Ghosh TS, O’Toole PW. The Healthy Microbiome—What Is the Definition of a Healthy Gut Microbiome? Gastroenterology. 2021;160(2):483–494. doi: 10.1053/j.gastro.2020.09.057

[2] Kinross JM, Darzi AW, Nicholson JK. Gut microbiome-host interactions in health and disease. Genome Medicine. 2011;3(3):14. doi: 10.1186/gm228

[3] Bidell MR, Hobbs AL V, Lodise TP. Gut microbiome health and dysbiosis: A clinical primer. Pharmacotherapy: The Journal of Human Pharmacology and Drug Therapy. 2022;42(11):849– 857. doi: 10.1002/phar.2731

[4] Nichols RG, Davenport ER. The relationship between the gut microbiome and host gene expression: a review. Human genetics. 2021;140(5):747–760. doi: 10.1007/s00439-020-02237-0

[5] Curry KD, Wang Q, Nute MG, Tyshaieva A, Reeves E, Soriano S, Wu Q, Graeber E, Finzer P, Mendling W, et al. Emu: species-level microbial community profiling of full-length 16S rRNA Oxford Nanopore sequencing data. Nature Methods. 2022;19(7):845–853. doi: 10.1038/s41592-022-01520-4

[6] Wilkins LJ, Monga M, Miller AW. Defining Dysbiosis for a Cluster of Chronic Diseases. Scientific Reports. 2019;9(1):12918.doi: 10.1038/s41598-019-49452-y

[7] Visconti A, Le Roy CI, Rosa F, Rossi N, Martin TC, Mohney RP, Li W, de Rinaldis E, Bell JT, Venter JC, et al. Interplay between the human gut microbiome and host metabolism. Nature Communications. 2019;10(1):4505. https://doi.org/10.1038/s41467-019-12476-z. doi: 10.1038/s41467-019-12476-z

[8] Schmidt TSB, Hayward MR, Coelho LP, Li SS, Costea PI, Voigt AY, Wirbel J, Maistrenko OM, Alves RJC, Bergsten E, et al. Extensive transmission of microbes along the gastrointestinal tract Garrett WS, Nieuwdorp M, Prodan A, O’Toole P, editors. eLife. 2019;8:e42693. doi: 10.7554/eLife.42693

[9] Park S-Y, Hwang B-O, Lim M, Ok S-H, Lee S-K, Chun K-S, Park K-K, Hu Y, Chung W-Y, Song N-Y. Oral-gut microbiome axis in gastrointestinal disease and cancer. Cancers (Basel). 2021;13(9):2124. doi: 10.3390/cancers13092124

[10] Solé C, Guilly S, Da Silva K, Llopis M, Le-Chatelier E, Huelin Patricia and Carol M, Moreira R, Fabrellas Núria and De Prada G, Napoleone L, Graupera Isabel and Pose E, et al. Alterations in gut microbiome in cirrhosis as assessed by quantitative metagenomics: Relationship with acute-on-chronic liver failure and prognosis. Gastroenterology. 2021;160(1):206–218.e13. doi: 10.1053/j.gastro.2020.08.054.

[11] Xu F, Fu Y, Sun T, Jiang Z, Miao Z, Shuai M, Gou W, Ling C, Yang J, Wang J, et al. The interplay between host genetics and the gut microbiome reveals common and distinct microbiome features for complex human diseases. Microbiome. 2020;8(1):145. doi: 10.1186/s40168-020-00923-9

[12] Wu G, Zhao N, Zhang C, Lam YY, Zhao L. Guild-based analysis for understanding gut microbiome in human health and diseases. Genome Medicine. 2021;13(1):22. doi: 10.1186/s13073-021-00840-y

[13] Dosajh A, Agrawal P, Chatterjee P, Priyakumar UD. Modern machine learning methods for protein property prediction. Current Opinion in Structural Biology. 2025;90:102990. doi: 10.1016/j.sbi.2025.102990

[14] Kanakala GC, Devata S, Chatterjee P, Priyakumar UD. Generative artificial intelligence for small molecule drug design. Current Opinion in Biotechnology. 2024;89:103175. doi: 10.1016/j.copbio.2024.103175

[15] Korlepara DB, C. S. V, Srivastava R, Pal PK, Raza SH, Kumar V, Pandit S, Nair AG, Pandey S, Sharma S, et al. PLAS-20k: Extended Dataset of Protein-Ligand Affinities from MD Simulations for Machine Learning Applications. Scientific Data. 2024;11(1):180. doi: 10.1038/s41597-023-02872-y

[16] Kiouri DP, Batsis GC, Mavromoustakos T, Giuliani A, Chasapis CT. Structure-Based Modeling of the Gut Bacteria–Host Interactome Through Statistical Analysis of Domain– Domain Associations Using Machine Learning. BioTech. 2025;14(1). doi: 10.3390/biotech14010013

[17] Curry KD, Nute MG, Treangen TJ. It takes guts to learn: machine learning techniques for disease detection from the gut microbiome. Emerging Topics in Life Sciences. 2021;5(6):815– 827. doi: 10.1042/ETLS20210213

[18] Giuffrè M, Moretti R, Tiribelli C. Gut microbes meet machine learning: the next step towards advancing our understanding of the gut microbiome in health and disease. International Journal of Molecular Sciences. 2023;24(6):5229. doi: 10.3390/ijms24065229

[19] Li P, Luo H, Ji B, Nielsen J. Machine learning for data integration in human gut microbiome. Microbial Cell Factories. 2022;21(1):241. doi: 10.1186/s12934-022-01973-4

[20] Asnicar F, Thomas AM, Passerini A, Waldron L, Segata N. Machine learning for microbiologists. Nature Reviews Microbiology. 2024;22(4):191–205. doi: 10.1038/s41579-023-00984-1

[21] Janiesch C, Zschech P, Heinrich K. Machine learning and deep learning. Electronic markets. 2021;31(3):685–695. doi: 10.1007/s12525-021-00475-2

[22] Dias R, Torkamani A. Artificial intelligence in clinical and genomic diagnostics. Genome Medicine. 2019;11(1):70. doi: 10.1186/s13073-019-0689-8

[23] Nethala TR, Sahoo BK, Srinivasulu P. Optimal gene therapy network: Enhancing cancer classification through advanced AI-driven gene expression analysis. e-Prime - Advances in Electrical Engineering, Electronics and Energy. 2024;7:100449. doi: 10.1016/j.prime.2024.100449

[24] Trebicka J, Bork P, Krag A, Arumugam M. Utilizing the gut microbiome in decompensated cirrhosis and acute-on-chronic liver failure. Nature Reviews Gastroenterology & Hepatology. 2021;18(3):167–180. doi: 10.1038/s41575-020-00376-3

[25] Oh M, Zhang L. DeepMicro: deep representation learning for disease prediction based on microbiome data. Scientific Reports. 2020;10(1):6026. doi: 10.1038/s41598-020-63159-5

[26] Syama K, Jothi JAA, Khanna N. Automatic disease prediction from human gut metagenomic data using boosting GraphSAGE. BMC Bioinformatics. 2023;24(1):126. doi: 10.1186/s12859-023-05251-x

[27] Chen X, Zhu Z, Zhang W, Wang Y, Wang F, Yang J, Wong K-C. Human disease prediction from microbiome data by multiple feature fusion and deep learning. iScience. 2022;25(4):104081. doi: 10.1016/j.isci.2022.104081

[28] Bang S, Yoo D, Kim S-J, Jhang S, Cho S, Kim H. Establishment and evaluation of prediction model for multiple disease classification based on gut microbial data. Scientific Reports. 2019;9(1):10189. doi: 10.1038/s41598-019-46249-x

[29] Olaguez-Gonzalez JM, Chairez I, Breton-Deval Luz and Alfaro-Ponce M. Machine learning algorithms applied to predict autism spectrum disorder based on gut microbiome composition. Biomedicines. 2023;11(10). doi: 10.3390/biomedicines11102633.

[30] Sharma D, Paterson AD, Xu W. TaxoNN: ensemble of neural networks on stratified microbiome data for disease prediction. Bioinformatics. 2020;36(17):4544–4550. doi: 10.1093/bioinformatics/btaa542

[31] Pietrucci D, Teofani A, Unida V, Cerroni R, Biocca S, Stefani A, Desideri A. Can gut Microbiota be a good predictor for Parkinson’s disease? A machine learning approach. Brain Sci. 2020;10(4):242. doi: 10.3390/brainsci10040242

[32] Li Z, Zhou J, Liang H, Ye L, Lan L, Lu F, Wang Q, Lei T, Yang X, Cui P, et al. Differences in alpha diversity of gut Microbiota in neurological diseases. Front. Neurosci. 2022;16:879318.

[33] Aryal S, Alimadadi A, Manandhar I, Joe B, Cheng X. Machine learning strategy for gut microbiome-based diagnostic screening of cardiovascular disease. Hypertension. 2020;76(5):1555–1562. doi: 10.3389/fnins.2022.879318

[34] Gupta VK, Kim M, Bakshi U, Cunningham KY, Davis JM, Lazaridis KN, Nelson H, Chia N, Sung J. A predictive index for health status using species-level gut microbiome profiling. Nature Communications. 2020;11(1):4635. doi: 10.1038/s41467-020-18476-8

[35] Reitmeier S, Kiessling S, Clavel T, List M, Almeida EL, Ghosh TS, Neuhaus K, Grallert H, Linseisen J, Skurk T, et al. Arrhythmic Gut Microbiome Signatures Predict Risk of Type 2 Diabetes. Cell Host & Microbe. 2020;28(2):258–272.e6. doi: 10.1016/j.chom.2020.06.004

[36] Sagi O, Rokach L. Ensemble learning: A survey. Wiley interdisciplinary reviews: data mining and knowledge discovery. 2018;8(4):e1249. doi: 10.1002/widm.1249

[37] Zhang J. Enhancing Medical Diagnostics with Machine Learning: A Study on Ensemb le Methods and Transfer Learning. Vol. 70. EDP Sciences; p. 02022. doi: 10.1051/itmconf/20257002022

[38] Nadarajah S, Mba JC, Rakotomarolahy P, Ratolojanahary HTJE. Ensemble Learning and an Adaptive Neuro-Fuzzy Inference System for Cryptocurrency Volatility Forecasting. Journal of Risk and Financial Management. 2025;18(2). doi: 10.3390/jrfm18020052

[39] Zhang Y, Liu J, Shen W. A Review of Ensemble Learning Algorithms Used in Remote Sensing Applications. Applied Sciences. 2022;12(17). doi: 10.3390/app12178654

[40] Aouedi O, Piamrat K, Parrein B. Ensemble-Based Deep Learning Model for Network Traffic Classification. IEEE Transactions on Network and Service Management. 2022;19(4):4124–4135. doi: 10.1109/TNSM.2022.3193748

[41] Mahajan P, Uddin S, Hajati F, Moni MA. Ensemble Learning for Disease Prediction: A Review. Healthcare. 2023;11(12). doi: 10.3390/healthcare11121808

[42] Yin X, Liu Q, Pan Y, Huang X, Wu J, Wang X. Strength of Stacking Technique of Ensemble Learning in Rockburst Prediction with Imbalanced Data: Comparison of Eight Single and Ensemble Models. Natural Resources Research. 2021;30(2):1795–1815. doi: 10.1007/s11053-020-09787-0

[43] Muller E, Algavi YM, Borenstein E. The gut microbiome-metabolome dataset collection: a curated resource for integrative meta-analysis. npj Biofilms and Microbiomes. 2022;8(1):79. doi: 10.1038/s41522-022-00345-5

[44] Pasolli E, Truong DT, Malik F, Waldron L, Segata N. Machine Learning Meta-analysis of Large Metagenomic Datasets: Tools and Biological Insights. PLoS Computational Biology. 2016;12(7):e1004977. doi: 10.1371/journal.pcbi.1004977

[45] Chawla N V, Bowyer KW, Hall LO, Kegelmeyer WP. SMOTE: synthetic minority over-sampling technique. Journal of artificial intelligence research. 2002;16:321–357. doi: 10.1613/jair.953

[46] Hasan BMS, Abdulazeez AM. A review of principal component analysis algorithm for dimensionality reduction. Journal of Soft Computing and Data Mining. 2021;2(1):20–30.

[47] Shetty SH, Shetty S, Singh C, Rao A. Supervised machine learning: algorithms and applications. Fundamentals and methods of machine and deep learning: algorithms, tools and applications. 2022:1–16. doi: 10.1002/9781119821908.ch1

[48] Eckhardt CM, Madjarova SJ, Williams RJ, Ollivier M, Karlsson J, Pareek A, Nwachukwu BU. Unsupervised machine learning methods and emerging applications in healthcare. Knee Surgery, Sports Traumatology, Arthroscopy. 2023;31(2):376–381. doi: 10.1007/s00167-022-07233-7

[49] Ladosz P, Weng L, Kim M, Oh H. Exploration in deep reinforcement learning: A survey. Information Fusion. 2022;85:1–22. doi: 10.1016/j.inffus.2022.03.003

