## Supplementary Word File containing the supporting information, including protocols, algorithms, methods, figures, and tables. for "Enhanced Prediction of Gut Microbiome–Related Diseases Using Hybrid Machine Learning Models"

**For**

**SI 1. Data pre-processing**

Table SI 1. The datasets and corresponding abbreviations

| Dataset Name | Year | Diseased Classes | Abbreviations | References |
| --- | --- | --- | --- | --- |
| Deep Micro |  | Obesity, Leanness | D1 | [1] |
|  | 2020 | Normal, T2D | D2 |  |
|  |  | Normal, IBD_crohn_disease, IBD_ulcerative_colitis | D3 |  |
|  |  | Normal, Cirrhosis | D4 |  |
|  |  | Normal, Cancer, Small Adenoma | D5 |  |
|  |  | Normal, T2D | D6 |  |
| Borenstein Lab | 2019 | History of Colorectal Surgery (HS), Normal, Multiple Polypoid, Stage 0, Stage I/II, Stage III,IV | Y1 | [2] |
| Kaggle | 2016 | Cancer, Cirrhosis, IBD_crohn_disease, IBD_ulcerative_colitis, Impaired_Glucose_Tolerance, Large_Adenoma, Leanness, Normal, Normal at risk, Obesity, Small_Adenoma, Stec2-Positive, T2D | K1 | [3] |
| TaxoNN | 2020 | Normal, Cirrhosis | T1 | [4] |
|  | 2020 | Normal, T2D | T2 |  |

Table SI 2. Original datasets dimensions before any pre-processing techniques.

| Diseased Categories (Abbreviations (Table SI 1)) | Number of Patients/ Sample size | Number of Features (Bacterial Species) |
| --- | --- | --- |
| D5 | 121 | 507 |
| D1 | 253 | 467 |
| D6 | 344 | 576 |
| D4 | 232 | 549 |
| D3 | 110 | 447 |
| D2 | 96 | 383 |
| Y1 | 347 | 16389 |
| K1 | 3072 | 3303 |
| T1 | 233 | 208 |
| T2 | 344 | 208 |

Table SI 3. The dimension of the datasets followed by cleaning, normalization and augmentation.

| Diseased Categories (Abbreviations (Table SI 1)) | Number of Patients/ Sample size | Number of Features (Bacterial Species) | Augmentation |
| --- | --- | --- | --- |
| D1 | 328 | 467 | Yes |
| D2 | 106 | 383 | Yes |
| D3 | 255 | 455 | Yes |
| D4 | 236 | 544 | Yes |
| D5 | 144 | 505 | Yes |
| D6 | 348 | 574 | Yes |
| Y1 | 762 | 16389 | Yes |
| K1 | 8054 | 3303 | Yes |
| T1 | 233 | 208 | No |
| T2 | 1740 | 208 | Yes |

**SI 2. ML based models**

Machine learning (ML) models are algorithms that learn patterns from data to make predictions or decisions without being explicitly programmed. These models are generally categorized into supervised learning (e.g., linear regression, support vector machines, random forests), unsupervised learning (e.g., k-means clustering, principal component analysis, DBSCAN), and reinforcement learning (e.g., Q-learning, deep Q-networks) techniques [5–7] In this context, supervised learning models are employed, as the datasets contain labelled target variables along with their corresponding ground truth values. These are used as baseline models in our current study, to compare the performances of the proposed hybrid models.

SI 2.1 Decision tree (DT)

DT consists of a tree-like structure that acts like a flowchart. The internal nodes are feature tests (or attribute tests), each branch is the result of that test, and each leaf node is a class label or decision. It is formed by recursively partitioning data based on feature values to maximize information gain or minimize impurity at every step.The model is interpretable and well-suited for both classification and regression tasks. Gini Impurity is a measure of the probability of incorrectly classifying a randomly chosen element from a dataset, which is calculated using (eqn. SI 1 )

Gini Impurity**:**$G=1-\sum_{i=1}^{k} p_{i}^{2}$	(SI 1)

Here, *K* is the number of classes, $p_{i}$ is the proportion of samples in class *i* at the node.

Weighted Gini Impurity: The weighted Gini impurity formula is used to measure the impurity of a decision tree node after a split, which is calculated using (eqn. SI 2)

$G_{split}= \frac{n_{l}}{n}G_{l}+\frac{n_{r}}{n}G_{r}$	(SI 2)

Here, $n_{l}, n_{r}$ are the number of samples in the left and right child nodes, n is the total number of samples in the parent node, and $G_{l}, G_{r}$ are the gini impurities of the left and right child nodes.

SI 2.2 Random forest (RF)

RF essentially is an ensemble of DT models [8]**,** whereby each individual models combine to raise accuracy and enhance robustness. The basis for such a combination depends on the fact that every tree differs in its construction since it is built for each dataset and a subset of features of the overall random feature. It generally doesn't overfit as long as more trees are incorporated. Classification uses majority votes in obtaining the dominance of the output predictions from each decision tree (100 in the present RF model), in contributing to the final estimated prediction aggregates.

In Figure SI 1. Random Forest, each decision tree differs from others because of the bootstrapping, that is, sample data are randomly selected and replaced at each split. As these trees train, having different subsets of data and looking at different features, this results in the building of different structures and decision boundaries.

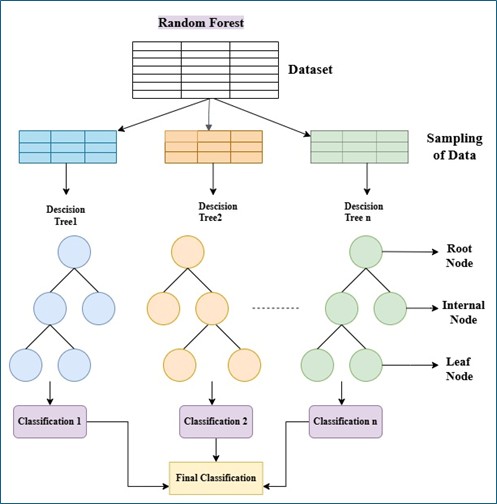

Figure SI 1. Random Forest model architecture.

SI 2.3 Logistic regression (LoR)

LoR is yet another linear model for binary classification problems. It calculates predictions of the likelihood that an input data point would fall into a particular class by feeding a weighted sum of its input features through the logistic or sigmoid function .[9] With the use of gradient descent and its variants, it is possible to find the best-fit coefficients for this model as well. LoR model works extremely well with linearly separable data.

z=$w^{'}x+b$	(SI 3)

y'=σ(z)	(SI 4)

Equations (SI 3) and (SI 4) represent the forward pass in a logistic regression model, where the input *x* is transformed using weights and biases (eqn. SI 3), and the sigmoid activation function (eqn. SI 4) maps the output to a probability for binary classification.

Where, z is the linear combination of weights and input features, w is the vector of weights, x is the vector of input features, b is the bias term, y' is the predicted output and σ(z) is the activation function.

SI 2.4 Support vector machine (SVM)

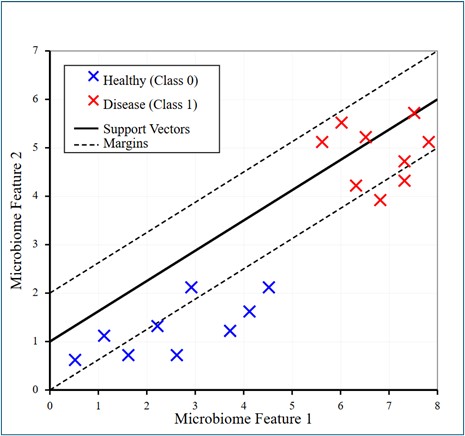

Figure SI 2. Schematic diagram for SVM model architecture.

SVM is a supervised ML model based on the maximum-margin approach between classes [10]. It separates points in space by creating a decision boundary using support vectors. SVM can be used for non-linear classification by making use of kernel functions (radial basis function kernel to handle non-linear decision boundaries, in the current model). This visualization of data in Figure SI 2 demonstrates how a Support Vector Machine (SVM) algorithm successfully separates healthy subjects from diseased subjects using two microbiome features. The scatter plot reveals a distinct pattern in which healthy subjects (blue X markers) are grouped in the lower-left area with smaller feature values, and diseased subjects (red X markers) are grouped in the upper-right area with larger feature values. The SVM has drawn an ideal decision boundary (solid black line) with optimal margins (dashed lines) defined by the support vectors on the critical. This separation in clear classes using the formula from (eqn. SI 5) implies that there is a high predictive value of the disease classification among these microbiome features.

$f\left( x \right)= w^{\tau}x+b$	(SI 5)

Here, $f\left( x \right)$ is the final prediction, *w* represents the weights, and *b* is the bias.

If $f\left( x \right)$ is greater than or equal to 0, classify *x* as class +1

If $f\left( x \right)$ is less than 0, classify *x* as class -1

SI 2.5 Naïve Bayes

Naive Bayes (NB) is a probabilistic classifier that assumes feature independence in a really strong manner.[11] Based on Bayes' theorem (reference), it computes the posterior probability for each class, given the feature values and predicts a class having the highest probability. Naive Bayes is a simple algorithm that works quite well on diverse tasks, due to it being efficient and effective with high-dimensional data. We have taken Gaussian NB classification function, used for continuous numerical values.

*P(y|x)*=$\frac{P(y)\prod_{i=1}^{n} P(X_{i}|y)}{P(x)}$	(SI 6)

In eqn. SI 6, *P_y_* is the prior probability of class y, $P(X_{i}|y)$ is the likelihood probability of the feature *X_i_* given class y and *P(x)* is the evidence (the total probability of observing the features, which is constant for all classes).

SI 2.6 Extreme gradient boost

Extreme gradient boost (XGB) is among the strongest and most general-purpose ML algorithms based on an enhanced version of gradient boosting.[12] It is usually used for classification and regression tasks. It builds decision trees sequentially, with each additional tree refining the shortcomings of the previous trees. As shown in Figure SI 3. XGB is different because it uses regularization (Lasso regressor (L1) and Ridge regressor (L2)), parallel processing, tree pruning, and missing value handling, all of which are faster and more accurate than traditional boosting methods. One can also tune some of the hyperparameters like the learning rate, number of trees, and depth for better control of accuracy, performance, and overfitting.

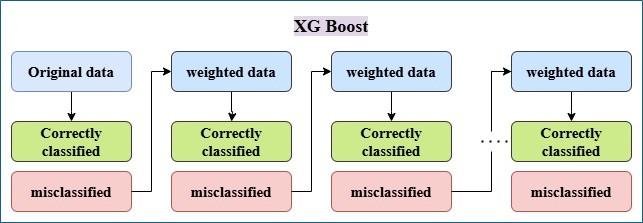

Figure SI 3. Extreme Gradient Boost (XGB) Architecture

SI 2.7 Multi-layer perceptron

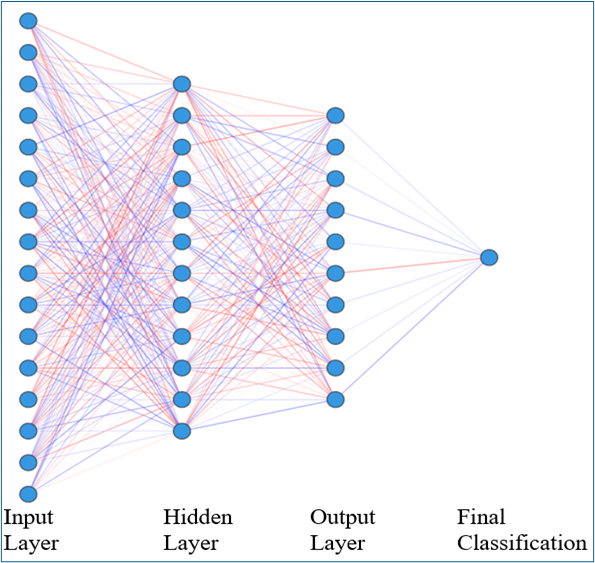

Figure SI 4. A common representation of NN architecture

A multilayer perceptron (MLP) is an NN that consists of several layers of nodes or neurons, one input layer, and either one or more hidden layers, finally followed by the output layer as shown in Figure SI 4. Each neuron of a given layer is connected to each neuron in the layer following, with their respective weights and biases tuned by backpropagation.[13] MLP, being an NN, is endowed with an activation function (ReLU or sigmoid), generally used to introduce nonlinearity and qualify it for more sophisticated functions. MLPs are nowadays used extensively in classification and regression problems where the relationship between features is nonlinear. We have used a simple architecture with one hidden layer with 100 neurons, with training for 1000 iterations, with “ReLu” as an activation function and “adam” as optimizer.

Input Layer:

$z^{1}=W^{1}x+b^{1},a^{1}=f^{1}\left( z^{1} \right)$	(SI 7)

$f^{l}(z)= max (0,z)$	(SI 8)

Hidden Layer:

$z^{l}=W^{l}a^{l-1}+b^{l},a^{l}=f^{l}\left( z^{l} \right)$	(SI 9)

Output Layer (classification):

$z^{l}=W^{l}a^{l-1}+b^{l},a^{l}=\sigma(z_{i})$	(SI 10)

Here, for the MLP, as shown in the Figure SI 4. l is the total number of layers including input and output layers, x is the input vector, $W^{l}$ is the weight matrix for layer l, $b^{l}$ is the bias vector for layer l, $z^{l}$ is the pre-activation values for layer l, $a^{l}$ is the post activation values for layer l, $f^{l}$ is the activation function for layer l (ReLu hidden layers) Forward Propagation of Neural Networks Equations (SI 7) to (SI 10) present the way in which information moves through the neural network. Wherein an activation function, beginning with the input layer transformation (eqn. SI 7), is followed by applying weights and biases (eqn. SI 8, most usually ReLU), and the hidden layers (eqn. SI 9) continue this process for the final output layer to use a softmax function for classification as shown in (eqn. SI 10). The size of the input layer would be the number of features that we are considering after PCA as it is dependent on the respective dataset. The hidden layer size is 100 neurons, and the output layer size is the number of unique classes respectively to the dataset.

SI 2.8 Neural Networks (NN)

NN are widely used for classification tasks, which capture complex patterns and hidden information. These consist of the input layer, a hidden layer, and an output layer.[14]

Here we have used the MLP architecture (see section SI 2.7) previously mentioned to train our model on the Kaggle Yachdia, TaxoNN (Cirrhosis and T2D) datasets (Table SI 1).

Additionally, we also implemented NN architectures for single/multi-class classification. It begins by one-hot encoding the labels and splitting the data into training and testing sets. A sequential neural network is created with two hidden layers (128 and 64 neurons, ReLU activation) and dropout layers (30%) to avoid overfitting. The model is compiled with the Adam optimizer and trained for 5 epochs.

Table SI 4. Summary of baseline models and configurations employed for classification.

| Model | Model Parameters |
| --- | --- |
| LR | penalty='l2', C=1.0 |
| NB | var_smoothing=1e-9 |
| SVM | RBF kernel |
| RF | n_estimators=100 |
| XGB | booster='gbtree', learning_rate=0.3, n_estimators=100 |
| MLP | 1 hidden layer (100 neurons), 1000 iterations, ReLU activation function, adam optimiser |
| NN | 2 hidden layers (128 and 64 neurons), ReLU activation function |

SI 2.9 Meta-classifiers in EM1 and EM2

Table SI 5. The hybrid models with the altered combinations of base models and metaclassifiers, and corresponding abbreviations.

| Hybrid Models | | Base Models | Meta classifiers |
| --- | --- | --- | --- |
| EM1 | | NB+XGB+SVM+RF | LoR |
| EM1 variations | EM1.1 | XGB+SVM+RF+LoR | NB |
|  | EM1.2 | NB+SVM+RF+LoR | XGB |
|  | EM1.3 | NB+XGB+RF+LoR | SVM |
|  | EM1.4 | NB+XGB+SVM+LoR | RF |
| EM2 | | RF+SVM | MLP |
| EM2 variations | EM2.1 | SVM+MLP | RF |
|  | EM2.2 | RF+MLP | SVM |

**SI 3. ML metrics**

The formulas for the respective metrics are given below in (eqns. SI 11 – SI 16).

Accuracy=$\frac{(TP+TN)}{(TP+TN+FP+FN)}$	(SI 11)

Precision=$\frac{(TP)}{(TP+FP)}$	(SI 12)

Recall=$\frac{(TP)}{(TP+FN)}$	(SI 13)

F1 Score=$\frac{(2\times P\times R)}{(P+R)}$	(SI 14)

Where, in (eqn. SI 11 – SI 14) TP is the number of True Positives, TN is the number of True Negatives, FP is the number of False Positives, and FN is the number of False Negatives.

AUC (Area under Curve):

Macro AUC=$\frac{1}{c}\sum_{c=1}^{c} {AUC}_{c}$	(SI 15)

Micro AUC=$\frac{\sum_{c=1}^{c} {TP}_{C}}{\sum_{C=1}^{C} ({TP}_{c}+{FP}_{C})}$	(SI 16)

In eqns. SI 15, 16, c is the number of classes. In our study, we have only used Micro AUC for model evaluation.

**SI 4. Complimentary Results**

Table SI 6. Tabulated Accuracies for the hybrid models with different combinations of meta-classifiers across the chosen datasets.

| Dataset | EM1 | EM1 variations | | | | EM2 | EM2 variations | |
| --- | --- | --- | --- | --- | --- | --- | --- | --- |
|  |  | EM1.1 | EM1.2 | EM1.3 | EM1.4 |  | EM2.1 | EM2.2 |
| D1 | 0.89 | 0.78 | 0.71 | 0.80 | 0.75 | 0.80 | 0.80 | 0.80 |
| D2 | 0.68 | 0.59 | 0.68 | 0.68 | 0.63 | 0.63 | 0.63 | 0.59 |
| D3 | 0.98 | 0.96 | 0.98 | 0.90 | 0.98 | 0.80 | 0.80 | 0.25 |
| D4 | 0.94 | 0.94 | 0.68 | 0.92 | 0.92 | 0.92 | 0.89 | 0.87 |
| D5 | 0.65 | 0.62 | 0.24 | 0.65 | 0.65 | 0.69 | 0.51 | 0.69 |
| D6 | 0.88 | 0.71 | 0.60 | 0.73 | 0.76 | 0.86 | 0.85 | 0.77 |
| K1 | 0.95 | 0.80 | 0.95 | 0.94 | 0.95 | 0.98 | 0.98 | 0.97 |
| Y1 | 0.92 | 0.92 | 0.90 | 0.91 | 0.92 | 0.92 | 0.92 | 0.92 |
| T1 | 0.89 | 0.89 | 0.80 | 0.89 | 0.87 | 0.91 | 0.87 | 0.89 |
| T2 | 0.98 | 0.97 | 0.97 | 0.93 | 0.97 | 0.98 | 0.98 | 0.97 |
| Average Accuracy | 0.87 | 0.82 | 0.75 | 0.83 | 0.84 | 0.84 | 0.82 | 0.77 |

Table SI 7. Performance measure with ensemble models before balancing the classes (data corresponding to Table SI 2). The data dimension for T1 was not augmented, and therefore not analysed as imbalanced classes, due to the class being already balanced.

| Datasets | Hybrid model | Accuracy | Precision | Recall | F1score | AUC | CV score  (Cross validation) |
| --- | --- | --- | --- | --- | --- | --- | --- |
| K1 | EM1 | 0.70 | 0.56 | 0.70 | 0.61 | 0.94 | 0.71 |
|  | EM2 | 0.70 | 0.62 | 0.70 | 0.65 | 0.95 | 0.72 |
| Y1 | EM1 | 0.43 | 0.07 | 0.16 | 0.10 | 0.71 | 0.34 |
|  | EM2 | 0.41 | 0.24 | 0.21 | 0.20 | 0.69 | 0.34 |
| T1 | EM1 | - | - | - | - | - | - |
|  | EM2 | - | - | - | - | - | - |
| T2 | EM1 | 0.60 | 0.61 | 0.60 | 0.60 | 0.68 | 0.65 |
|  | EM2 | 0.65 | 0.65 | 0.65 | 0.65 | 0.70 | 0.59 |

Table SI 8. Performance measure with ensemble models, after balancing the classes (T1 is already balanced: Table SI 2). “*Reference*” here denotes the AUC from the base papers.

| Datasets | Hybrid model | Accuracy | Precision | Recall | F1score | AUC | CV score | AUC (*Reference*) |
| --- | --- | --- | --- | --- | --- | --- | --- | --- |
| K1 | EM1 | 0.95 | 0.95 | 0.95 | 0.95 | 0.99 | 0.94 | 0.96 |
|  | EM2 | 0.98 | 0.98 | 0.98 | 0.98 | 0.99 | 0.97 |  |
| Y1 | EM1 | 0.92 | 0.93 | 0.92 | 0.92 | 0.98 | 0.89 | 0.85 |
|  | EM2 | 0.92 | 0.93 | 0.93 | 0.93 | 0.99 | 0.89 |  |
| T1 | EM1 | 0.89 | 0.89 | 0.89 | 0.89 | 0.95 | 0.85 | 0.93 |
|  | EM2 | 0.91 | 0.92 | 0.91 | 0.92 | 0.92 | 0.83 |  |
| T2 | EM1 | 0.98 | 0.98 | 0.98 | 0.98 | 0.99 | 0.95 | 0.73 |
|  | EM2 | 0.98 | 0.98 | 0.98 | 0.98 | 0.99 | 0.95 |  |

Table SI 9. Classwise F1 score for the ensemble models.

| Dataset Name | Diseased Categories (Abbreviations) | Classes | F1 score | |
| --- | --- | --- | --- | --- |
|  |  |  | EM1 | EM2 |
| Deep Micro | D1 | Obesity | 0.89 | 0.79 |
|  |  | Leaness | 0.90 | 0.81 |
|  | D2 | Normal | 0.59 | 0.64 |
|  |  | T2D | 0.74 | 0.64 |
|  | D3 | Normal | 0.97 | 0.94 |
|  |  | IBD_crohn_disease | 0.99 | 0.99 |
|  |  | IBD_ulcerative_colitis | 0.97 | 0.94 |
|  | D4 | Normal | 0.93 | 0.91 |
|  |  | Cirrhosis | 0.94 | 0.92 |
|  | D5 | Normal | 0.55 | 0.64 |
|  |  | Cancer | 0.70 | 0.60 |
|  |  | Small Adenoma | 0.75 | 0.92 |
|  | D6 | Normal | 0.89 | 0.86 |
|  |  | T2D | 0.89 | 0.85 |
| Borenstein Lab | Y1 | Normal | 0.84 | 0.87 |
|  |  | History of Colorectal Surgery (HS) | 0.95 | 0.93 |
|  |  | Multiple Polypoid | 0.99 | 0.99 |
|  |  | Stage 0 | 0.96 | 0.99 |
|  |  | Stage I/II | 0.84 | 0.84 |
|  |  | Stage III,IV | 0.97 | 0.97 |
| Kaggle | K1 | Normal | 0.78 | 0.82 |
|  |  | Normal at risk | 0.99 | 0.99 |
|  |  | Cancer | 0.99 | 0.99 |
|  |  | Cirrhosis | 0.94 | 0.97 |
|  |  | IBD_crohn_disease | 0.99 | 0.99 |
|  |  | IBD_ulcerative_colitis | 0.95 | 0.97 |
|  |  | Impaired_Glucose_Tolerance | 0.99 | 0.99 |
|  |  | Large_Adenoma | 0.99 | 0.99 |
|  |  | Leanness | 0.96 | 0.97 |
|  |  | Obesity | 0.90 | 0.90 |
|  |  | Small_Adenoma | 0.97 | 0.99 |
|  |  | Stec2-Positive | 0.99 | 0.99 |
|  |  | T2D | 0.89 | 0.92 |
| TaxoNN | T1 | Normal | 0.88 | 0.90 |
|  |  | Cirrhosis | 0.91 | 0.92 |
|  | T2 | Normal | 0.98 | 0.99 |
|  |  | T2D | 0.98 | 0.98 |

**SI 5. References**

[1]	Oh M, Zhang L. DeepMicro: deep representation learning for disease prediction based on microbiome data. Scientific Reports. 2020;10(1):6026. doi: 10.1038/s41598-020-63159-5

[2]	Muller E, Algavi YM, Borenstein E. The gut microbiome-metabolome dataset collection: a curated resource for integrative meta-analysis. npj Biofilms and Microbiomes. 2022;8(1):79. doi: 10.1038/s41522-022-00345-5

[3]	Pasolli E, Truong DT, Malik F, Waldron L, Segata N. Machine Learning Meta-analysis of Large Metagenomic Datasets: Tools and Biological Insights. PLoS Computational Biology. 2016;12(7):e1004977. doi: 10.1371/journal.pcbi.1004977

[4]	Sharma D, Paterson AD, Xu W. TaxoNN: ensemble of neural networks on stratified microbiome data for disease prediction. Bioinformatics. 2020;36(17):4544–4550. doi: 10.1093/bioinformatics/btaa542

[5]	Shetty SH, Shetty S, Singh C, Rao A. Supervised machine learning: algorithms and applications. Fundamentals and methods of machine and deep learning: algorithms, tools and applications. 2022:1–16. doi: 10.1002/9781119821908.ch1

[6]	Eckhardt CM, Madjarova SJ, Williams RJ, Ollivier M, Karlsson J, Pareek A, Nwachukwu BU. Unsupervised machine learning methods and emerging applications in healthcare. Knee Surgery, Sports Traumatology, Arthroscopy. 2023;31(2):376–381. doi: 10.1007/s00167-022-07233-7

[7]	Ladosz P, Weng L, Kim M, Oh H. Exploration in deep reinforcement learning: A survey. Information Fusion. 2022;85:1–22. doi: 10.1016/j.inffus.2022.03.003

[8]	Biau G, Scornet E. A random forest guided tour. TEST. 2016;25(2):197–227. doi: 10.1007/s11749-016-0481-7

[9]	Wright RE. Logistic regression. In: Reading and understanding multivariate statistics. Washington,  DC,  US: American Psychological Association; 1995. p. 217–244. doi: 10.4236/ojl.2021.102010

[10]	Hearst MA, Dumais ST, Osuna E, Platt J, Scholkopf B. Support vector machines. IEEE Intelligent Systems and their Applications. 1998;13(4):18–28. doi: 10.1109/5254.708428

[11]	Rish I. An empirical study of the naive Bayes classifier. In: IJCAI 2001 workshop on empirical methods in artificial intelligence. Vol. 3. Seattle, WA, USA; 2001. p. 41–46. doi: 10.4236/jsea.2023.166009

[12]	Santhanam R, Uzir N, Raman S, Banerjee S. Experimenting XGBoost Algorithm for Prediction and Classification of Different Datasets. 2017. doi: 10.54097/2dq7z209

[13]	Pinkus A. Approximation theory of the MLP model in neural networks. Acta Numerica. 1999;8:143–195. doi: 10.1017/S0962492900002919

[14]	Gu J, Wang Z, Kuen J, Ma L, Shahroudy A, Shuai B, Liu T, Wang X, Wang G, Cai J, et al. Recent advances in convolutional neural networks. Pattern Recognition. 2018;77:354–377. doi: 10.1016/j.patcog.2017.10.013
